# Systematic analysis of IL-1 cytokine signaling by high-risk HPV oncoproteins

**DOI:** 10.1101/2022.08.25.505364

**Authors:** Paola Castagnino, Hee Won Kim, Long Kwan Metthew Lam, Devraj Basu, Elizabeth A. White

## Abstract

Human papillomavirus (HPV) E6 and E7 oncoproteins are expressed at all stages of HPV-mediated carcinogenesis and are essential drivers of cancers caused by high-risk HPV. Some of the activities of HPV E6 and E7, such as their interactions with host cellular tumor suppressors, have been characterized extensively. There is less information about how high-risk HPV E6 and E7 alter cellular responses to cytokines that are present in HPV-infected tissues and are an important component of the tumor microenvironment. We used several models of HPV oncoprotein activity to assess how E6 and E7 alter the cellular response to the pro-inflammatory cytokine IL-1β. Models of early-stage HPV infection and of established HPV-positive head and neck cancers exhibited similar dysregulation of IL-1 pathway genes and suppressed transcriptional responses to IL-1β treatment. Such overlap in cell responses supports that changes induced by HPV E6 and E7 early in infection could persist and contribute to a dysregulated immune environment throughout carcinogenesis. HPV E6 and E7 also drove the upregulation of several suppressors of IL-1 cytokine signaling, including SIGIRR, both in primary keratinocytes and in cancer cells. SIGIRR knockout was insufficient to increase IL-1β-dependent gene expression in the presence of HPV16 E6 and E7, suggesting that multiple suppressors of IL-1 signaling contribute to dampened IL-1 responses in HPV16-positive cells.

**IMPORTANCE:** Human papillomavirus (HPV) infection is responsible for nearly 5% of the worldwide cancer burden. HPV-positive tumors develop over years to decades in tissues that are subject to frequent stimulation by pro-inflammatory cytokines. However, the effects of HPV oncoproteins on the cellular response to cytokine stimulation are not well defined. We analyzed IL-1 cytokine signaling in several models of HPV biology and disease. We found that HPV16 E6 and E7 oncoproteins mediate a broad and potent suppression of cellular responses to IL-1β in models of both early and late stages of carcinogenesis. Our data provide a resource for future investigation of IL-1 signaling in HPV-positive cells and cancers.

## INTRODUCTION

High-risk HPV cause infections in squamous epithelial tissues that can persist and sometimes develop into cancers of the cervix, oropharynx, and other mucosa. The HPV E6 and E7 oncoproteins are expressed during initial HPV infection and are the major drivers of malignant progression in HPV-infected cells. HPV E6 and E7 bind to host cellular targets to reprogram infected cells, in doing so establishing an altered cell state that enables virus replication. High-risk HPV E7 bind and degrade the retinoblastoma tumor suppressor (RB1), releasing E2F transcription factors and allowing passage through the G1/S checkpoint (1-3). High-risk HPV E6 proteins bind the cellular ubiquitin ligase E6AP to form a complex that targets p53 for degradation, thereby blocking apoptotic signaling that would otherwise be triggered by E7 (4, 5). High-risk HPV E7 proteins also bind and degrade the tumor suppressor PTPN14, activating the YAP1 oncoprotein (6-8). Together E6 and E7 enable HPV-infected cells to persist in the basal layer of stratified epithelia and promote cellular reprogramming in differentiated cells. Over 50% of cervical cancers and 90% of HPV-positive head and neck cancers are caused by HPV16.

Only a small fraction of HPV infections lead to precancerous lesions and cancers, indicating that additional cell-intrinsic and exogenous influences contribute to HPV-related malignant progression (9). Genetic factors, tobacco and alcohol use, and other environmental influences may cooperate with the growth-promoting activities of HPV oncoproteins to enable carcinogenic progression. Among environmental factors, inflammation and the presence of pro-inflammatory cytokines are key features of the tumor microenvironment that can support carcinogenesis (10, 11). IL-1β is a pro-inflammatory cytokine that has been studied extensively in a variety of disease states. It is a master regulator upstream of TNFα, IL-6, and many other components of inflammatory signaling (12). Like other pro-inflammatory cytokines, IL-1β has both pro- and anti-tumorigenic effects that vary by cell and tumor type, but the balance of evidence supports a pro-tumorigenic role. HPV infect keratinocytes in stratified epithelia. Keratinocytes are important components of barrier immune responses that both produce and respond to pro-inflammatory cytokines. Keratinocytes initially respond to stimulation with IL-1β by expression of genes related to epidermal development, mitosis, and interferon receptors and subsequently by upregulation of cytokines and chemokines in an NF-κB and MyD88-dependent manner (13).

Some aspects of IL-1 signaling in HPV-infected cells have been characterized. Using cell-based models, several groups have characterized IL-1β secretion from keratinocytes expressing HPV E6 and/or E7. In general, HPV-immortalized or transformed cells are impaired in IL-1β secretion compared to normal cells (14) and HPV-negative oral cancer cell lines secrete more IL-1β than HPV-positive oral cancer cell lines (15). A few publications have proposed mechanisms by which HPV oncoproteins suppress IL-1β production. Niebler and colleagues found that HPV16 E6 decreases IL-1β protein levels by recruiting E6AP to target pro-IL-1β for proteasome-mediated degradation (16). Ainouze and colleagues implicated HPV16 E6 in suppressing IL-1β, but via a transcriptional mechanism dependent on downregulation of IRF6 (17).

However, there is less information about how cells expressing HPV E6 and E7 respond to paracrine IL-1β stimulation. The impact of high-risk HPV oncoproteins on the expression of cellular genes involved in IL-1 family cytokine responses is not well documented and the studies on how HPV-positive cells produce IL-1β did not characterize how HPV-positive cancer cells respond to IL-1β. There is also no consensus on whether HPV oncoproteins dysregulate IL-1 signaling in similar ways both in early stages of infection and later in disease progression. Here, we undertook an unbiased analysis of IL-1 signaling in several models of HPV-mediated disease. We observed that genes associated with IL-1 sensing are dysregulated in the presence of high-risk HPV E6 and E7 in models of both early infection and cancer. Consistent with a downregulation of pro-inflammatory cytokines and upregulation of certain IL-1 inhibitors in the presence of E6 and E7, there was broad global suppression of the cellular response to IL-1β in early infection and disease models. Although one inhibitor of IL-1 signaling, SIGIRR, was consistently upregulated in the presence of high-risk HPV E6 and E7, SIGIRR knockout was insufficient to restore normal IL-1 signaling in HPV-positive cells. Our studies provide global data sets that will inform future research on how pro-inflammatory cytokine signaling is altered in HPV-infected cells.

## RESULTS

### HPV16 oncoproteins alter the expression of IL-1 family cytokines and IL-1 regulatory genes

To make a comprehensive assessment of the effects of HPV16 oncoproteins on gene expression related to IL-1 signaling, we analyzed RNA-seq data from primary human foreskin keratinocytes (HFK) expressing HPV16 E6 and HPV16 E7 (HFK-HPV16 E6/E7). We selected 21 genes that represent core components of IL-1 signaling: nine IL-1 related cytokines, seven receptors and coreceptors, and five negative regulators or other inhibitors (Figure 1A). We assessed the expression of each gene in our RNA-seq data, finding that most cytokine gene expression and some receptor gene expression decreased in HFK-HPV16 E6/E7. In contrast, some of the negative regulators of IL-1 signaling were upregulated in the presence of HPV oncoproteins in one or more replicate cell lines. *SIGIRR*, which encodes a negative regulator of IL-1 receptor and Toll-like receptor signaling (18), was consistently upregulated in HFK-HPV16 E6/E7 cells compared to matched HFK-GFP controls. We used qRT-PCR and Western blot to validate levels of HPV16 E6, HPV16 E7, and *SIGIRR* RNAs and SIGIRR protein in the HFK cell lines (Figure 1B, 1C), finding that SIGIRR RNA and protein levels are elevated in the presence of HPV16 E6 and E7. To determine whether upregulation of *SIGIRR* was conserved among multiple HPV genotypes, we generated HFK that stably express E6 and E7 from HPV16, HPV18, and HPV6, as well as matched vector controls. We found that SIGIRR RNA and protein levels were increased in the presence of high-risk HPV16 and HPV18 oncoproteins, but not in cells expressing low-risk HPV6 E6 and E7 (Supplemental Figure 1). Consistent with a previous report, we observed that pro-IL-1β protein levels were reduced in HFK-HPV16 E6/E7 compared to controls (16) but found that other HPV E6/E7 did not lower pro-IL-1β levels.

**Figure 1.**
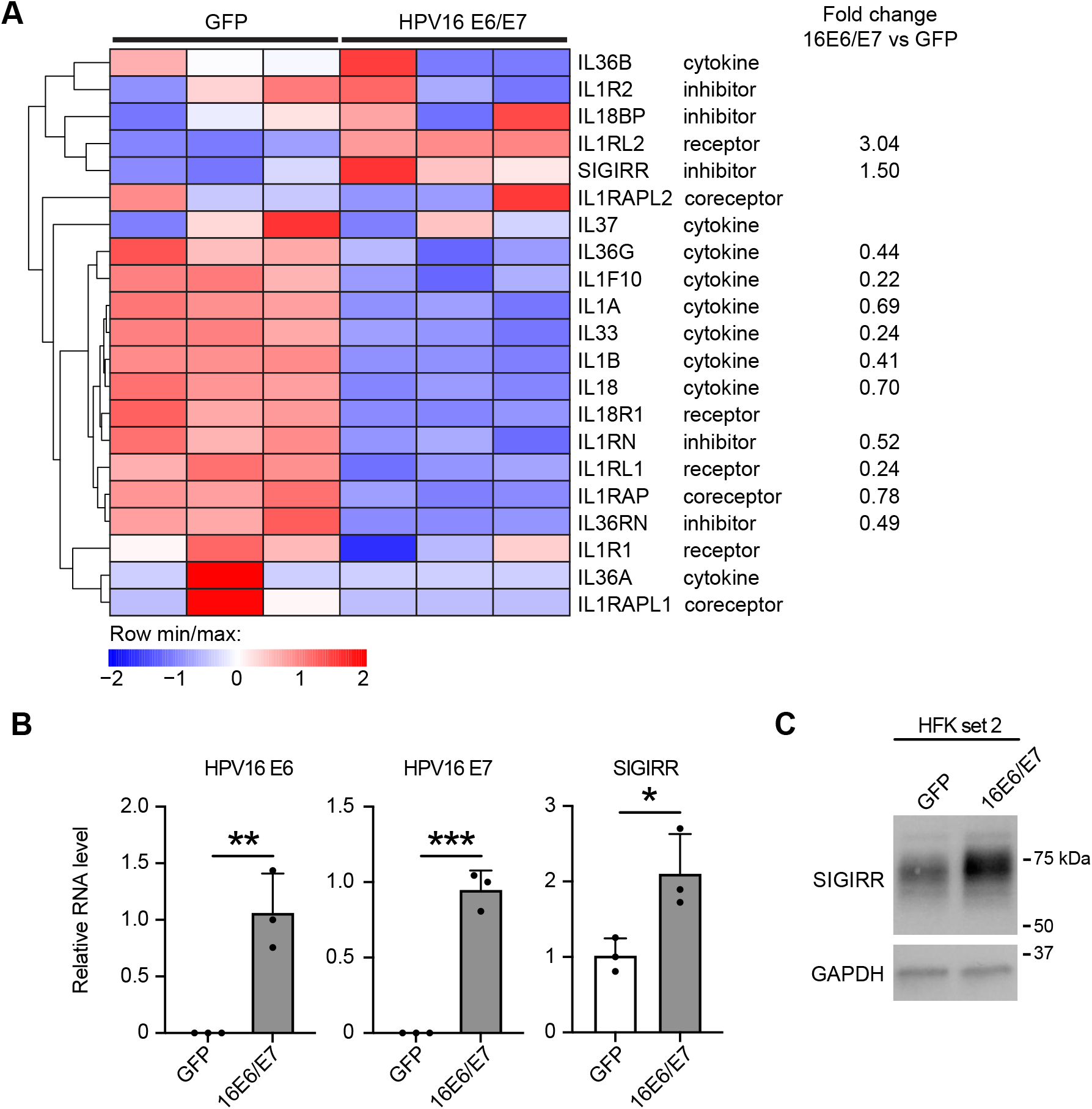
HPV16 oncoproteins dysregulate expression of genes involved in IL-1 signaling. A) Primary human foreskin keratinocytes (HFK) were transduced with retroviruses encoding HPV16 E6 and HPV16 E7 or with matched GFP control retroviruses and selected with appropriate antibiotics. RNA-seq and bioinformatic analysis was performed on polyA-selected RNA from triplicate cell lines. Heat map and fold change values indicate differential expression of IL-1 pathway cytokines and regulatory proteins. B) RNA from the same cell lines was analyzed by qRT-PCR using primers specific for HPV16 E6, HPV16 E7, or SIGIRR. Graphs display mean ± SD of the relative RNA level (vs GAPDH for HPV16 E6 and HPV16 E7, vs GAPDH for SIGIRR), dots represent individual cell lines. Statistical significance was assessed using an unpaired t-test, *, p<0.05; **, p<0.01; ***, p<0.001. C) Protein lysates from two of the six cell lines in (A) and (B) were separated by SDS-PAGE and analyzed in Western blots with antibodies to SIGIRR and GAPDH.

### IL-1 family cytokine and IL-1 regulatory gene expression differ in HPV-negative versus HPV-positive cancers

Having determined that high-risk HPV oncoproteins alter the expression of genes related to IL-1 signaling in primary HFK, we next sought to determine whether there is differential expression of the same IL-1 pathway genes in HPV-positive versus HPV-negative cancers. We focused on oropharyngeal squamous cell carcinomas (OPSCC), a subset of head and neck squamous cell carcinomas (HNSCC), which can be caused by HPV infection (HPV-positive) or be of non-viral etiology (HPV-negative). Most HPV-positive OPSCC are caused by HPV16 (19). Using RNA-seq data from 28 HPV-negative and 53 HPV-positive OPSCC in The Cancer Genome Atlas (20-22), we observed that certain IL-1 pathway genes were differentially expressed in HPV-negative vs. HPV-positive cancers (Figure 2A). Several genes followed the same expression pattern observed in HFK: *IL1A, IL1B*, and *IL36RN* levels were lower in HPV-positive OPSCC than in HPV-negative OPSCC (Figure 2B, 2C). In contrast, *SIGIRR* expression was higher in HPV-positive OPSCC than in HPV-negative samples. There were minimal differences in mutation or amplification rates in the sequenced genomes of the same OPSCC samples (Supplemental Figure 2), with a trend towards more frequent mutation of *IL18BP* in HPV-negative OPSCC. Finally, we tested whether patterns of differential gene expression in HPV-negative versus HPV-positive cancers were reflected in a panel of patient-derived xenograft (PDX) models established from 11 HPV-negative and 8 HPV-positive human HNSCC (Figure 3, Supplemental Figure 3, Supplemental Table 1). The trends of lower *IL1A, IL1B*, and *IL36RN* expression and higher *SIGIRR* expression in HPV-positive compared to HPV-negative samples were similar in the PDX samples to the trends that we observed in the gene expression data from TCGA.

**Figure 2.**
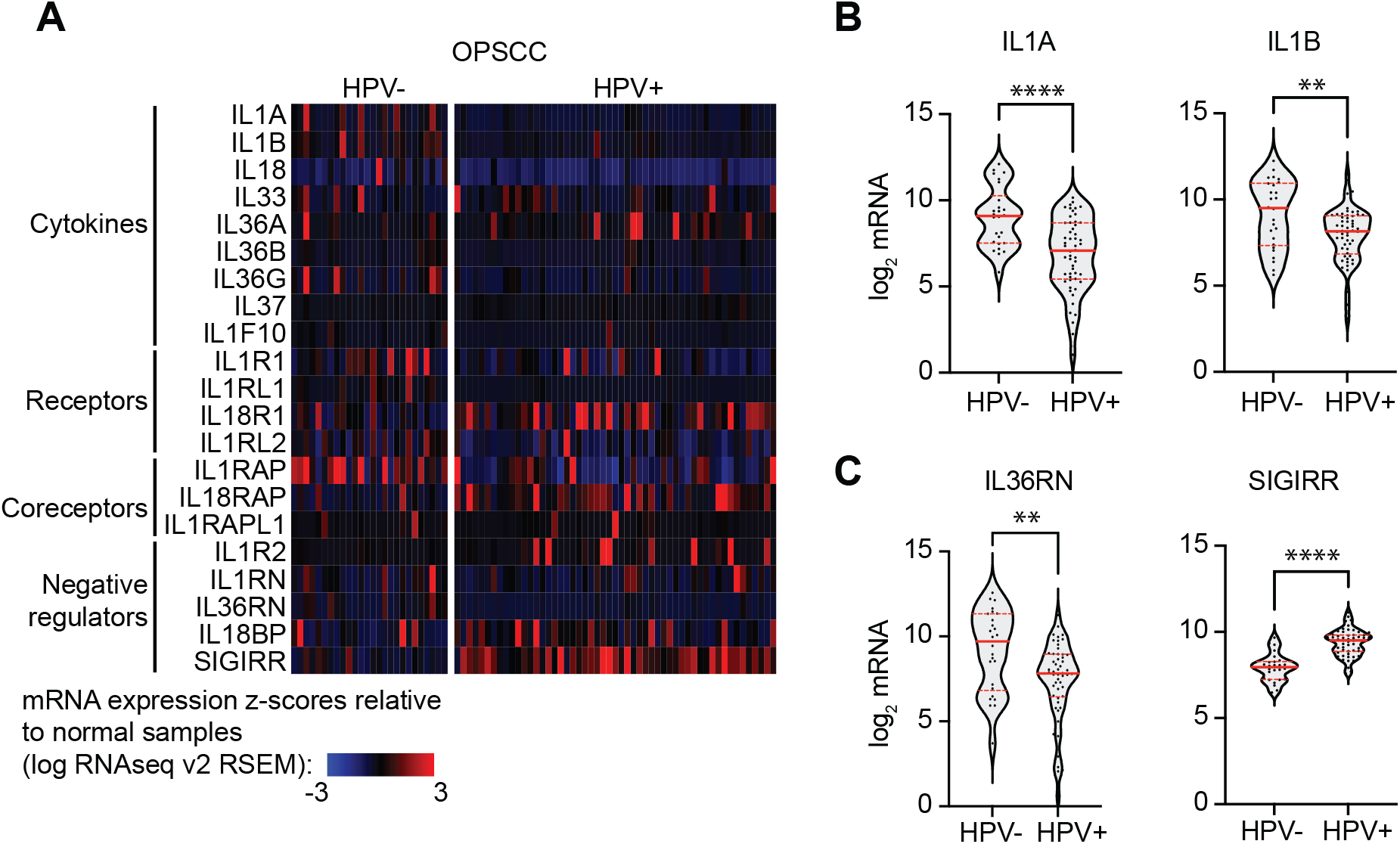
Genes related to IL-1 signaling are differentially expressed in HPV-positive versus HPV-negative oropharyngeal squamous cell carcinomas (OPSCC). Gene expression data from 28 HPV-negative and 53 HPV-positive OPSCC in The Cancer Genome Atlas (TCGA) was accessed using cBioPortal. A) Heat map displays summary mRNA expression data for selected genes encoding IL-1 related cytokines and IL-1 regulatory proteins. B-C) Violin plots display individual sample data from the same OPSCC for B) selected differentially expressed cytokines and C) selected differentially expressed negative regulators. Solid horizontal bar indicates median, dashed horizontal bars indicate top and bottom quartiles. Statistical significance was assessed using an unpaired t-test with Welch’s correction. ***, p<0.001; ****, p<0.0001.

**Figure 3.**
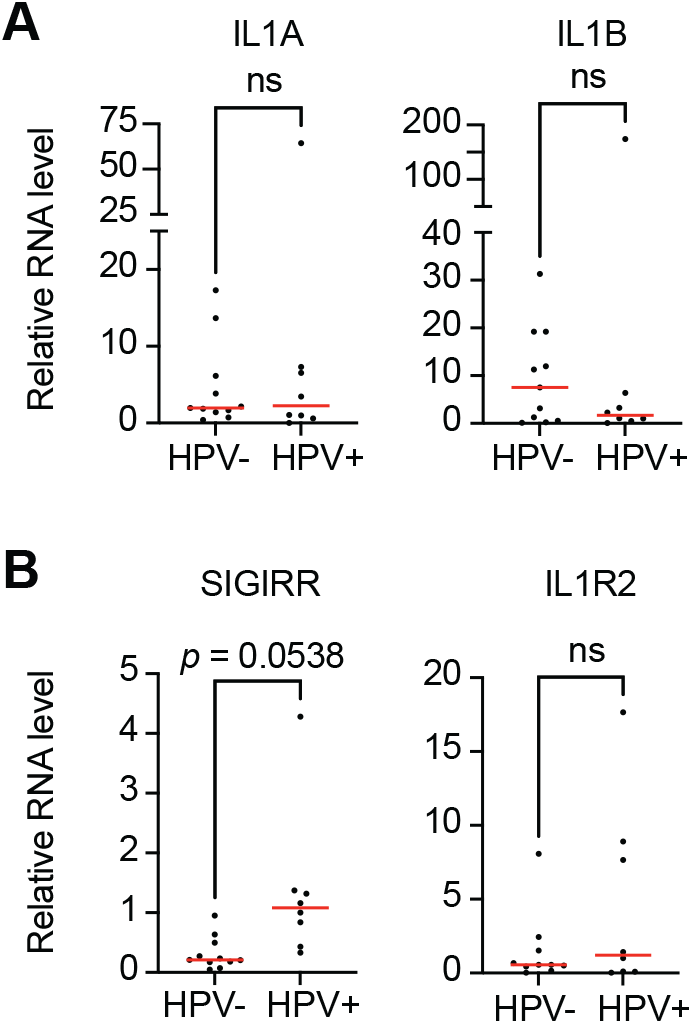
Differential expression of IL-1 regulatory proteins in HPV-negative and HPV-positive patient-derived xenografts. Total RNA was purified from 11 HPV-negative and 8 HPV-positive patient-derived xenografts (PDX) and qRT-PCR was used to assess gene expression of A) IL1A and IL1B or B) negative regulators of IL-1 signaling SIGIRR and IL1R2. Data is expressed relative to G6PD expression. Statistical significance was determined using unpaired t test with Welch’s correction. p-values below 0.1 are indicated. ns, not significant.

Both HFK-based models of HPV oncoprotein activity and data from patient samples reflect differential expression of genes related to IL-1 signaling in HPV-positive versus HPV-negative samples. Next, we tested whether the same gene expression differences were exhibited by a panel of established HNSCC cell lines. We measured several IL-1 related transcripts in two HPV-negative (SCC4, SCC15) and four HPV-positive (SCC47, SCC90, SCC152, VU147T) HNSCC cell lines, finding that SCC47 cells express more *IL1A and IL1B* than the other cell lines (Figure 4A). There was a trend towards higher expression of IL-1 negative regulators *IL1R2* and *SIGIRR* in three of the four HPV-positive cell lines (Figure 4B). Unlike in the HFK-based models of HPV oncoprotein activity, elevated levels of *SIGIRR* transcript in the HNSCC cell lines did not appear to correlate with differences in SIGIRR protein levels (Figure 4C).

**Figure 4.**
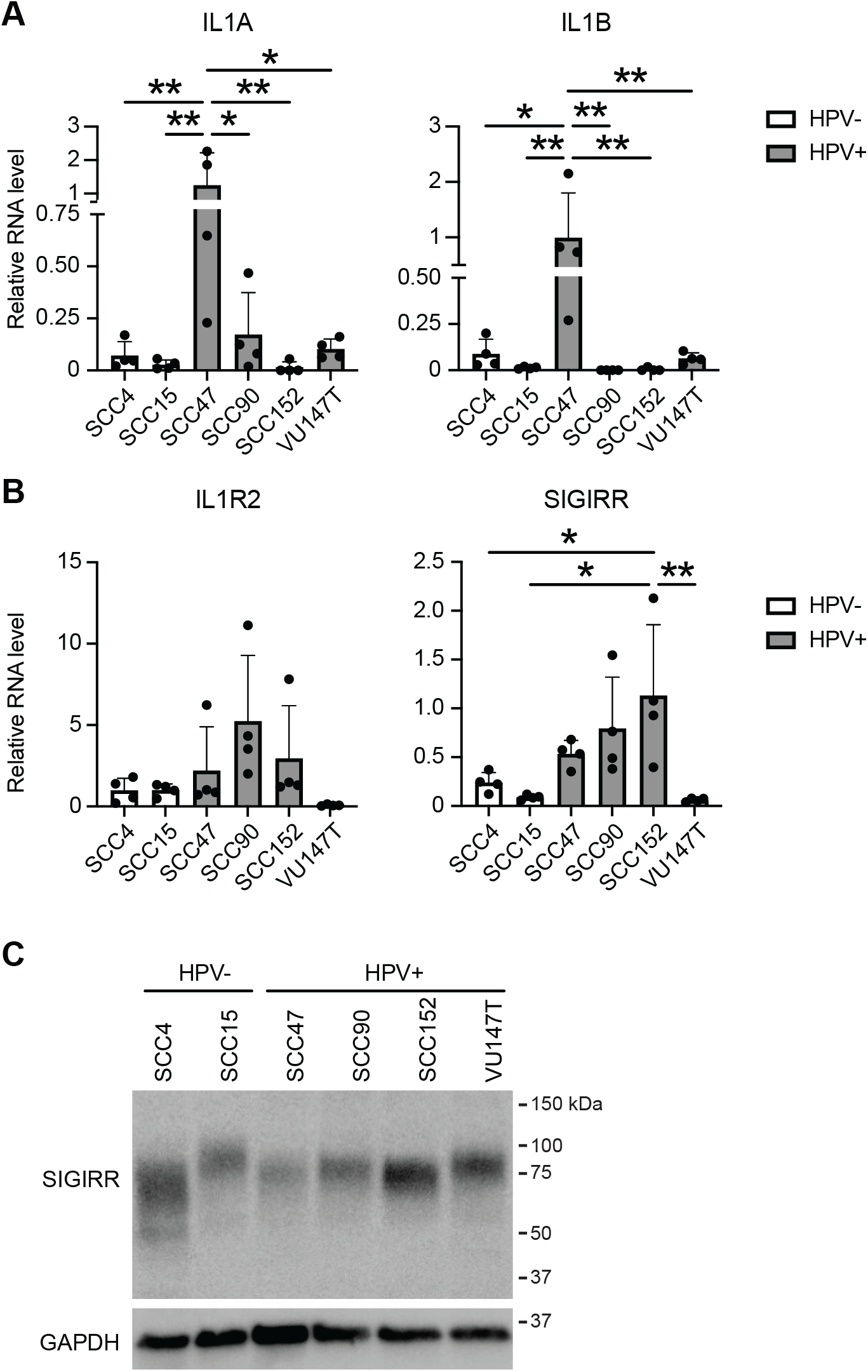
Differential expression of IL-1 family cytokines and IL-1 regulatory proteins in HPV-negative and HPV-positive HNSCC cell lines. Total RNA was purified from four passages each of two HPV-negative (SCC4, SCC15) and four HPV-positive (SCC47, SCC90, SCC152, VU147T) HNSCC cell lines. qRT-PCR was used to assess gene expression of A) IL1A and IL1B or B) selected negative regulators of IL-1 signaling. Data is expressed relative to G6PD expression. Statistical significance was determined using ordinary one-way ANOVA with Sidak’s multiple comparisons test. *, p<0.05; **, p<0.01. C) Protein lysates from the cell lines in (A-B) were separated by SDS-PAGE and analyzed in Western blots with antibodies to SIGIRR and GAPDH.

### HPV16 oncoproteins suppress the transcriptional response to IL-1β in human keratinocytes

Based on previous reports and our observations that IL-1 related genes are differentially expressed in models of HPV oncoprotein activity, we speculated that HPV16 E6 and E7 might mediate large-scale changes in the cellular response to IL-1β. We treated three independent HFK-16E6/E7 cell lines and three matched HFK-GFP controls with IL-1β and performed RNA-seq to assess global transcriptional changes (Figure 5, Supplemental Tables 2-5). In HFK-GFP there was a rapid and pronounced response to IL-1β treatment. Nearly all the genes differentially expressed upon IL-1β treatment were upregulated, and pathway analysis revealed that as expected, most upregulated genes were associated with Gene Ontology (GO) terms related to cellular inflammatory responses (Figure 5A, Supplemental Table 4). In comparison, the transcriptional response to IL-1β in HFK-16E6/E7 was suppressed. The genes that were most highly upregulated by IL-1β treatment in HFK-GFP were induced minimally or not at all by IL-1β treatment in HFK-16E6/E7 (Figure 5A). Although cells responded to IL-1β both in the presence and absence of HPV16 oncoproteins, HPV16 E6 and HPV16 E7 reduced both the number of genes induced by cytokine treatment and the magnitude of induction of IL-1 responsive genes (Figure 5A, 5B). Analysis of genes associated with the GO term ‘Response to Interleukin 1’ revealed a similar trend (Supplemental Figure 4). The genes that were associated with the response to IL-1 and that were most induced by IL-1β in HFK-GFP were not induced in HFK-16E6/E7. Cytokine treatment did not significantly alter the ability of HPV16 E6/E7 to promote expression of genes involved in DNA replication or to dysregulate the expression of genes related to keratinocyte differentiation (Figure 5C). We conclude that HPV16 oncoproteins enable a global suppression of the transcriptional response to IL-1β in human keratinocytes.

**Figure 5.**
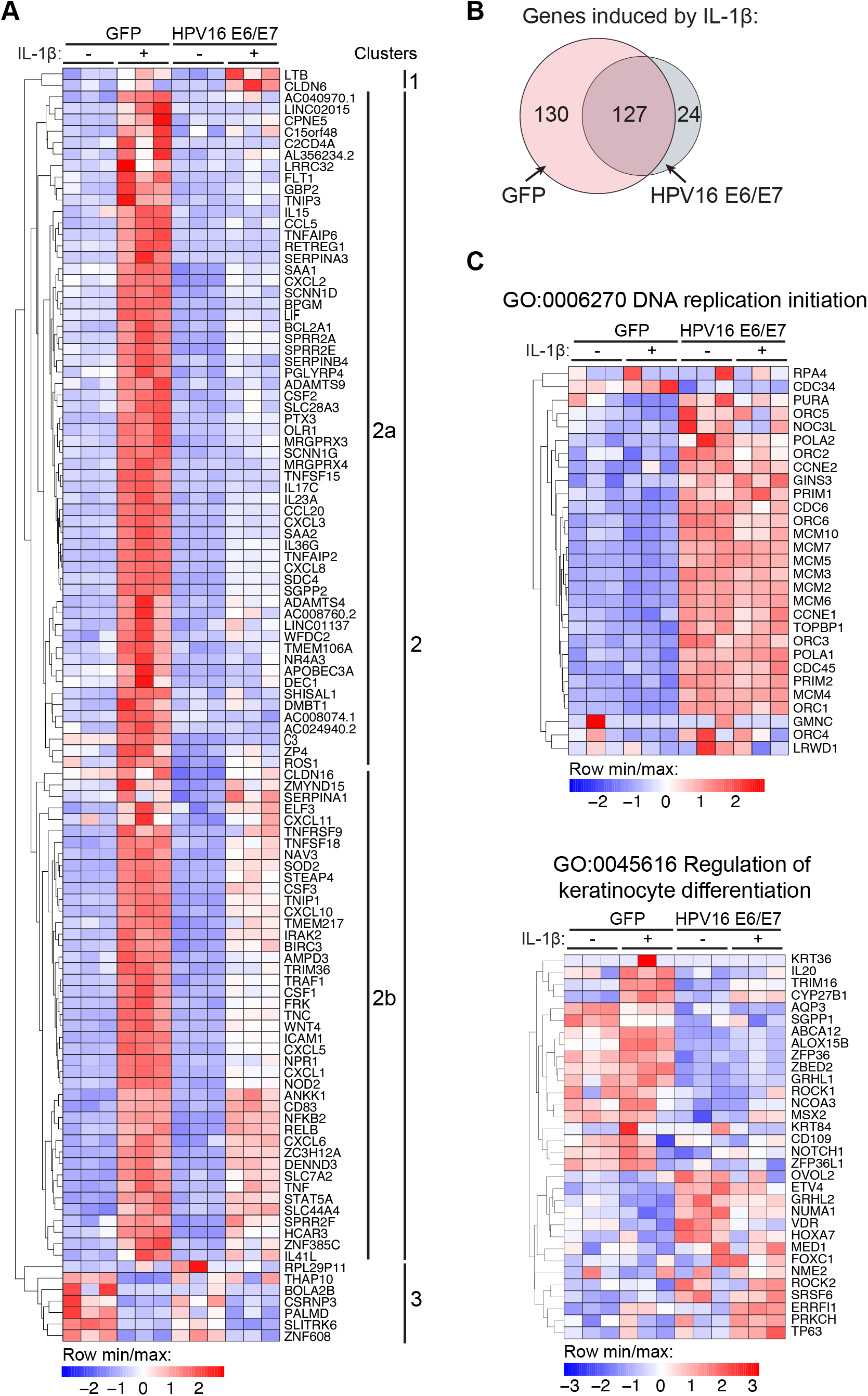
HPV16 oncoproteins suppress the transcriptional response to IL-1β. Primary HFK stably expressing HPV16 E6 and HPV16 E7 or matched GFP control cells were treated with 0.5 ng/ml IL-1β for 3h or left untreated. RNA-seq and bioinformatic analysis was performed on polyA-selected RNA from triplicate cell populations. A) Heat map displays expression values for genes that are differentially expressed by ≥2.5-fold in GFP cells treated with IL-1β compared to untreated GFP cells. Gene Ontology analysis was performed on clusters of differentially expressed genes using DAVID. B) Venn diagram indicates overlap between genes upregulated ≥‘1.5-fold upon IL-1β treatment in GFP cells or HPV16E6/E7 cells. C) Heat maps display expression data for genes associated with selected GO terms.

### HPV16 oncoproteins relieve IL-1β-associated growth inhibition in human keratinocytes

The observation that HPV16 E6/E7 had a broad and potent repressive effect on IL-1-associated gene expression led us to test whether the presence of HPV oncoproteins altered cell growth in the presence of IL-1β. We grew triplicate populations of HFK-HPV16E6/E7 and HFK-GFP in the presence and absence of IL-1β for approximately one month (Figure 6). HFK-GFP grew in culture for a few passages, but soon exhibited a decreased replicative capacity associated with cellular senescence. Culture in the presence of IL-1β further slowed the growth of HFK-GFP. In contrast, HPV16 E6/E7 conferred a growth advantage on cells that was equally pronounced in the presence and absence of IL-1β. We conclude that a growth-suppressive effect of IL-1β on human keratinocytes is alleviated in the presence of HPV16 oncoproteins.

**Figure 6.**
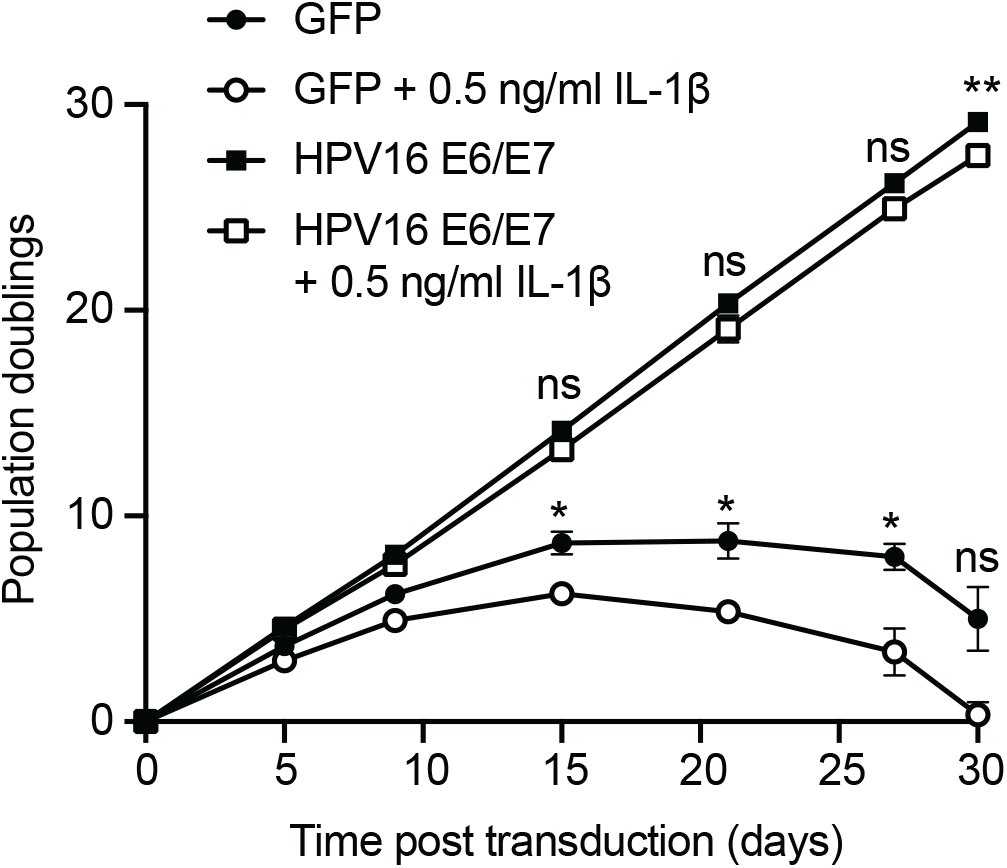
HPV16 E6/E7 alleviate the growth-suppressive effect of IL-1β on primary keratinocytes. Primary HFK were transduced with retroviruses encoding HPV16 E6 and E7 or matched GFP controls. Each cell population was cultured with or without 0.5 ng/ml IL-1β for 30 days and population doublings were tracked with each passage. Graph displays the mean ± SD of three replicate cell populations per condition. Statistical significance was determined by two-way ANOVA with Tukey’s multiple comparisons test (ns, not significant; *,p<0.05; **,p<0.01).

### HPV-positive HNSCC cell lines are less responsive to IL-1β than HPV-negative HNSCC cell lines

IL-1 pathway genes were suppressed in the presence of HPV16 E6/E7 in HFK and in cancer models, and IL-1β responsive gene expression was suppressed in HFK-16E6/E7. We next tested whether the cellular response to IL-1β was suppressed in HNSCC cell lines. We selected several IL-1β responsive genes from the RNA-seq data (Figure 4) and measured their expression in two HPV-negative and four HPV-positive HNSCC cell lines in the presence and absence of IL-1β (Figure 7). *CXCL8* is a canonical IL-1β responsive gene that was induced several-fold in HPV-negative SCC4 cells and was modestly induced by IL-1β treatment in HPV-negative SCC15 and HPV-positive SCC47 cells (Figure 7A). In three of the four HPV-positive SCC cell lines (SCC90, SCC152, and VU147T), *CXCL8* levels were low in untreated cells and exhibited minimal induction upon IL-1β treatment. Similarly, *CLXCL1* was strongly induced by IL-1β treatment in HFK-GFP cells. In both HPV-negative SCC cell lines, *CXCL1* was expressed in untreated cells and was induced several-fold upon IL-1β treatment (Figure 7B). In contrast, in the four HPV-positive cell lines, *CXCL1* expression was low in untreated cells and was not induced upon IL-1β treatment. Another gene, *SDC4*, that was IL-1β-inducible in HFK-GFP cells, was detectable but not significantly induced by IL-1β treatment in most HPV-positive and HPV-negative SCC cell lines (Figure 7C).

**Figure 7.**
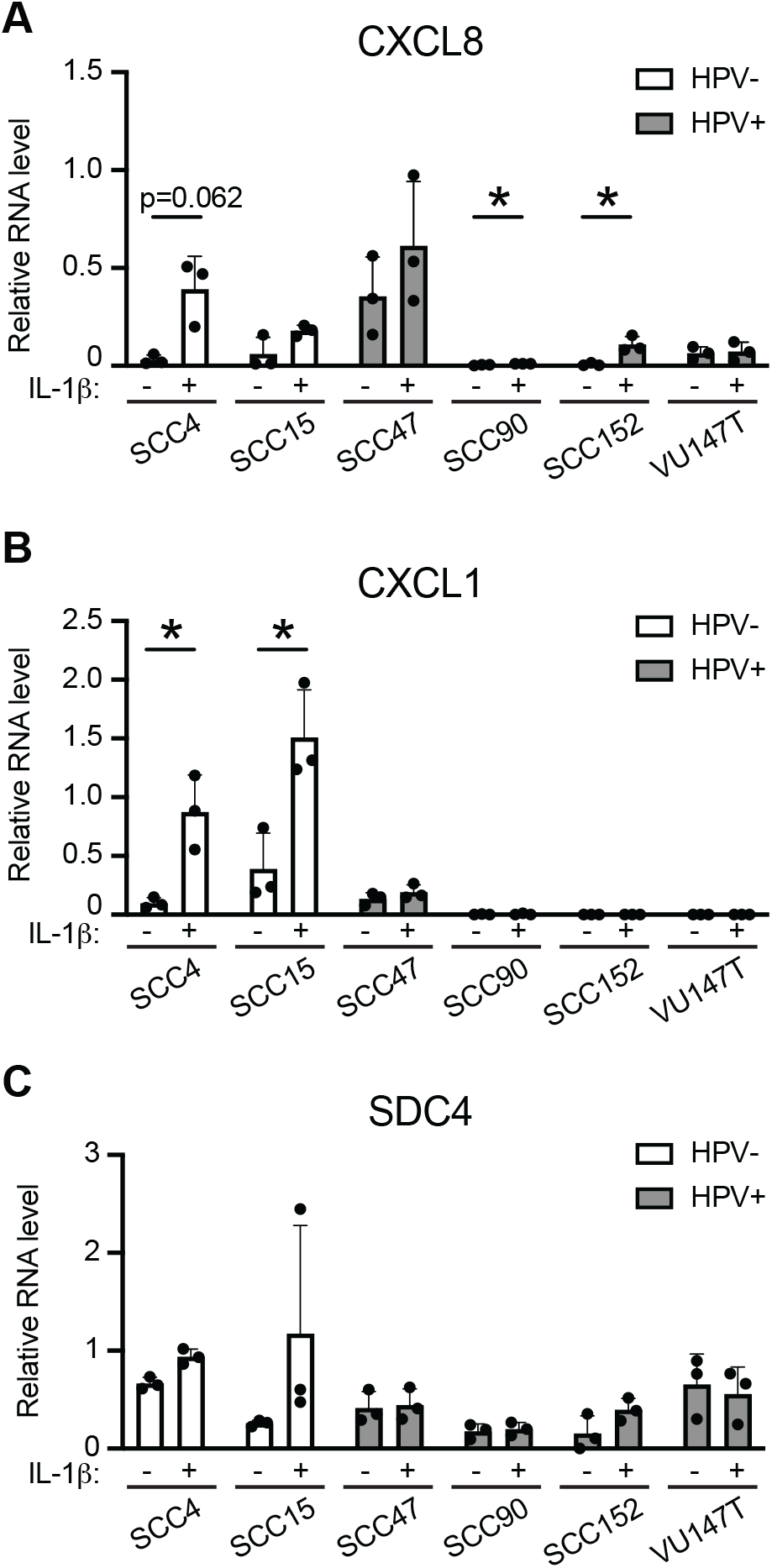
HPV-negative HNSCC cell lines are more responsive to IL-1β treatment than HPV-positive HNSCC cell lines. Two HPV-negative (SCC4, SCC15) and four HPV-positive (SCC47, SCC90, SCC152, VU147T) HNSCC cell lines were treated with 0.5 ng/ml IL-1β for 3h or left untreated. qRT-PCR was used to assess gene expression of A) CXCL8, B) CXCL1, or C) SDC4. Data is expressed relative to G6PD (for CXCL8) or GAPDH (for CXCL1 and SDC4) expression. Statistical significance was determined by unpaired t-test (*,p<0.05 or as indicated).

### *SIGIRR* is not responsible for the suppressed transcriptional response to IL-1β in the presence of HPV16 E6/E7

The consistent dampening of the response to IL-1β and upregulation of the IL-1 inhibitor SIGIRR in the presence of HPV16 E6/E7 led us to test whether SIGIRR might contribute to suppression of the IL-1 response in HPV-positive cells. We generated SIGIRR knockout cells in HFK-16E6/E7 (Figure 8A) and tested their response to IL-1β, finding that CXCL8 and CXCL1 were equally induced by IL-1β in the presence and absence of SIGIRR (Figure 8B). Similarly, knockout of SIGIRR from two different HPV16-positive HSNCC cell lines (Figure 8C) did not increase the transcriptional response of the cells to IL-1β (Figure 8D). We conclude that the IL-1 suppressive effect of HPV16 E6/E7 is not solely dependent on SIGIRR in HFK or in human cancer cell lines.

**Figure 8.**
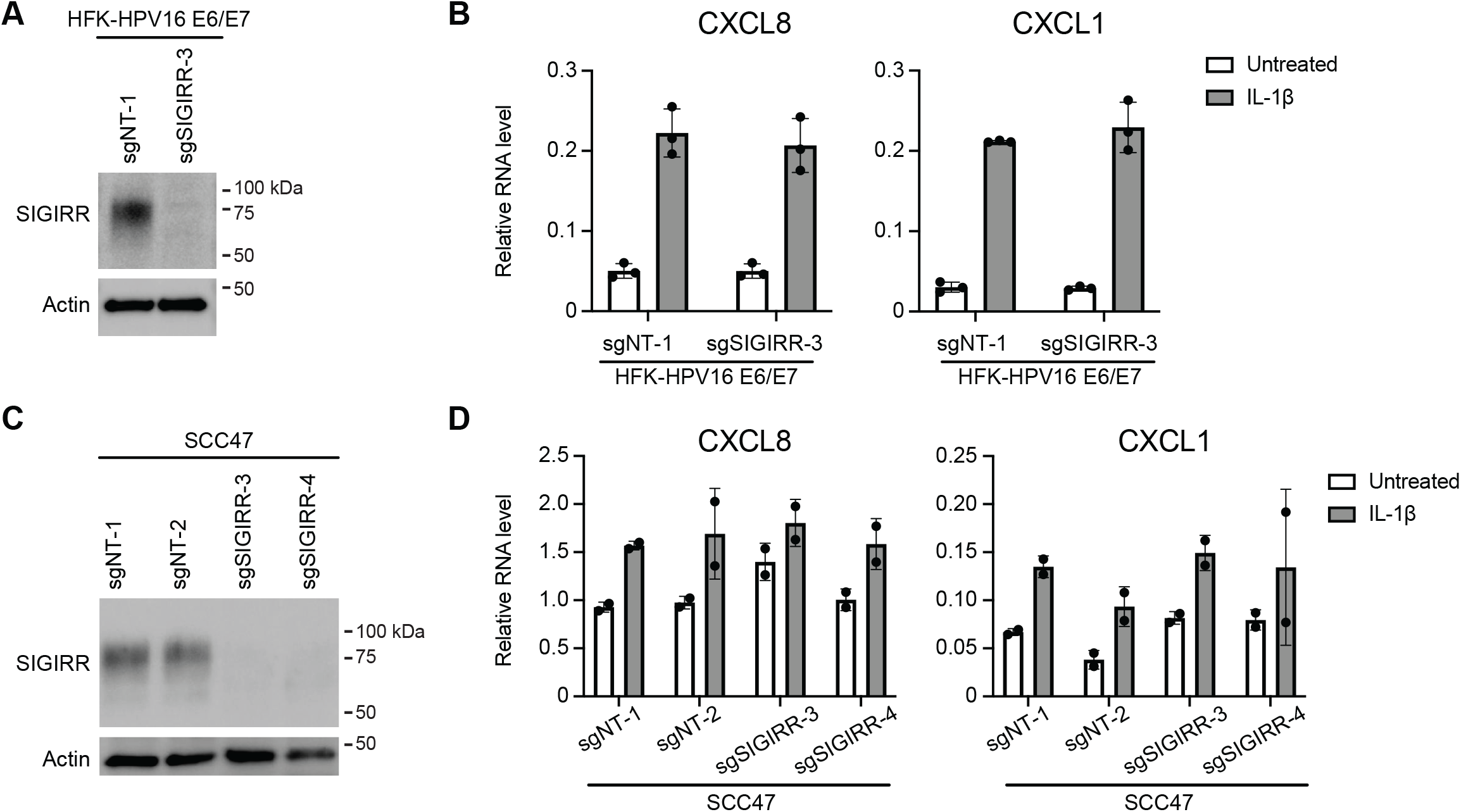
SIGIRR is not the primary driver of the suppressed response to IL-1β in cells expressing HPV16 E6/E7. A) HFK expressing HPV16 E6 and E7 were transduced with LentiCRISPRv2 vectors encoding a non-targeting (NT-2) sgRNA or an sgRNA targeting SIGIRR. SIGIRR depletion was confirmed by Western blot. B) Cells were treated with 0.5 ng/ml IL-1β for 3h or left untreated. qRT-PCR was used to assess gene expression of CXCL8 or CXCL1. Data is expressed relative to GAPDH expression. C) HPV-positive SCC47 cells were transduced with LentiCRISPRv2 vectors encoding non-targeting (NT) sgRNAs or sgRNAs targeting SIGIRR. SIGIRR depletion was confirmed by Western blot. D) Cells were treated with 0.5 ng/ml IL-1β for 3h or left untreated. qRT-PCR was used to assess gene expression of CXCL8 or CXCL1. Data is expressed relative to GAPDH expression.

## DISCUSSION

Initial infection of a keratinocyte by HPV is often said to provoke a minimal immune response. Even so, most HPV infections are cleared by the immune system. Some differences in immune cell recruitment to HPV-positive lesions and cancers have been characterized. Regressing cervical lesions are infiltrated by CD4+ and CD8+ T cells and macrophages whereas persistent HPV-positive lesions that progress to cancer are infiltrated by cancer-promoting tumor associated macrophages and depleted of protective Langerhans cells (23-27). Transcriptional data from HPV-positive vs HPV-negative head and neck squamous cell carcinomas (HNSCC) in The Cancer Genome Atlas indicates that HPV-positive tumors are more inflamed than their HPV-negative counterparts in terms of chemokines and of markers of T-cell recruitment (28). However, not all immune cell populations are enriched in HPV-positive tumors. Al-Sahaf and colleagues determined that HPV-negative tumors contain more neutrophils and express more of the neutrophil chemoattractant CXCL8 compared to HPV-positive tumors (15, 29). Overall, it is apparent that immune cell recruitment differs in HPV-positive vs HPV-negative tumors and that chronic inflammation could contribute to HPV-mediated carcinogenesis (30, 31).

Here, we undertook an unbiased analysis of IL-1 signaling in several models of HPV oncoprotein activity and HPV-associated disease. A global analysis of gene expression in keratinocytes expressing HPV16 E6 and E7 oncoproteins revealed overall downregulation of expression of many IL-1 family cytokines and an upregulation of certain negative regulators of IL-1 signaling including IL18BP and SIGIRR (Figure 1). Some aspects of this phenotype, particularly the downregulation of cytokine expression and upregulation of *SIGIRR*, were also observed in gene expression data from human OPSCC and HNSCC, in patient-derived xenograft models of HPV disease, and in HPV-positive HSNCC cell lines Figures 2-4). Notably, in the human cancer, PDX, and cancer cell line experiments, certain aspects of dysregulated IL-1 signaling were specific to HPV-positive, but not HPV-negative, cancer models. Continuing our experiments, we conducted a global analysis of gene expression in the presence and absence of IL-1β in HFK expressing HPV16 E6 and E7. We found that the presence of E6 and E7 was associated with an overall suppression of the keratinocyte response to IL-1β (Figure 5). IL-1β treatment did not affect the ability of HPV oncoproteins to promote cell cycle-related gene expression or to dysregulate keratinocyte differentiation. IL-1β had a growth-suppressive effect on HFK that was alleviated in the presence of HPV16 E6 and E7 (Figure 6). Findings in cancer cell lines were like those from the keratinocyte models, with several HPV-positive cancer cell lines exhibiting a suppressed response to IL-1β whereas HPV-negative cancer cells still responded to cytokine treatment (Figure 7).

Our findings are consistent with other data on differential cytokine signaling in HPV-transformed vs normal cells. HPV-positive cells generally produce lower levels of IL-1 cytokines compared to other cells (14). There is less information about how HPV-positive cells respond to or grow in the presence of IL-1 family cytokines, as might occur in the HPV-positive tumor microenvironment. Our observations that both basal IL-1 family gene expression and the response to IL-1β are reduced in HPV oncogene expressing cells may implicate a master regulator of gene expression sich as NF-κB. HPV-encoded proteins have several effects on NF-κB signaling, including that HPV E7 can inhibit NF-κB activity dependent on amino acids in the E7 C-terminus (32, 33). Future experiments could aim to further test whether effects of HPV16 E7 and/or E7 on NF-κB signaling account for the suppressed response to IL-1β we observed here.

In addition, our finding that certain inhibitors of IL-1 familty cytokine signaling are upregulated by HPV oncoproteins are consistent with other reports. For instance, IL18BP expression was increased in keratinocytes expressing high- and low-risk HPV E7 that were treated with IFN-γ (34). Both *IL18BP* and *SIGIRR*, another negative regulator of IL-1 signaling, were upregulated in our RNAseq data. Although SIGIRR was consistently upregulated in the presence of high-risk HPV E6 and E7, SIGIRR knockout did not increase IL-1β-responsive gene expression in E6/E7-expressing cells (Figure 8). We hypothesize that differential expression of multiple components of the IL-1 pathway contributes to the suppressed response in the presence of HPV oncoproteins. We also note that SIGIRR has been implicated in suppressing the response to other cytokines including IL-37 and to TLR-dependent responses (18, 35-41). Future experiments will aim to determine the consequence of SIGIRR upregulation in HPV-positive cells, in particular whether increased *SIGIRR* expression causes HPV-infected cells to mount a dampened transcriptional response to other IL-1 family cytokines or to stimuli of TLR-dependent signaling. Additional existing observations support the importance of interactions between HPV-infected cells and IL-1 family cytokines. Certain genetic polymorphisms in *IL1RA* are associated with decreased risk of developing cervical cancer and there are reports of different polymorphisms in *IL1B* that are either protective against or confer risk of developing cervical cancer (42-45). Some of these sequence changes have not been well characterized with respect to their activating or inhibitory effects on IL-1β secretion or activity.

Motivation for understanding how HPV oncoproteins influence IL-1 signaling extends beyond the study of HPV disease alone. HPV-positive cancers and non-cancerous oral lesions occur more frequently in people living with HIV, even those on antiretroviral therapy (PLWH/ART) than in HIV-uninfected populations (46-58). IL-1β is one of several pro-inflammatory cytokines that are consistently significantly upregulated in circulation and/or in tissues in HIV+ individuals on ART (59-66). There is a major gap in knowledge regarding why PLWH/ART exhibit increased rates of HPV-associated disease. A better understanding of how HPV-positive cells proliferate in the presence of pro-inflammatory cytokines will help to address the question of whether HPV E6 and E7 promote tumorgenesis in the HIV-infected tumor microenvironment.

## MATERIALS AND METHODS

### Plasmids and cloning

The HPV16 E7 ORF was cloned into MSCV-neo C-V5 GAW destination vector using Gateway recombination. The remaining MSCV-GFP, 16E6, and empty vector controls have been previously described (8, 67, 68). LentiCRISPRv2 vectors were cloned according to standard protocols using sgRNA sequences as contained in the Broad Institute Brunello library (69). A complete list of all plasmids used in this study is in Supplemental Table 6.

### Cell culture

Deidentified primary human foreskin keratinocytes (HFK) were provided by the University of Pennsylvania Skin Biology and Disease Resource-Based Center (SBDRC). HFK were cultured in K-SFM (Invitrogen) as previously described (70). Population doublings were calculated based upon the number of cells collected and re-plated at each passage. HNSCC cell lines were cultured in DMEM/F-12 (Invitrogen) containing 10% Fetal Bovine Serum and 50 ug/mL Gentamicin. For IL-1β treatment, recombinant human IL-1β was diluted in cell culture media at a final concentration of 0.5 ng/ml and cells were refed with the mixture for 3h or as indicated in the text. Retroviruses and lentiviruses were generated and used to established stable cell lines as previously described (67).

### Western blotting

Cells were lysed in RIPA buffer with phosphatase and protease inhibitors [150 mM sodium chloride, 5 mM EDTA, 50 mM Tris pH 7.8, 1% NP-40, 0.5% sodium deoxycholate, 0.1% SDS, 25 mM sodium fluoride, 0.1 mM sodium orthovanadate, 5 mM β-glycerophosphate, 1 mM phenylmethylsulfonyl fluoride, 5 μM leupeptin, and 1.4 μM pepstatin A] and lysates were separated on Tris/Glycine SDS-PAGE gels and transferred to polyvinylidene difluoride (PVDF). Membranes were blocked in Tris-buffered saline containing 5% nonfat dried milk and 0.05% Tween 20 (TBS-T), then incubated with primary antibodies that detect SIGIRR (Santa Cruz Biotechnology sc-271864), pro-IL-1β (R&D MAB201), GAPDH (Invitrogen MA5-15738), and Actin (Cell Signaling Technology 3700). After washing with TBS-T, membranes were incubated with horseradish peroxidase-coupled secondary antibodies and imaged using chemiluminescent substrate on an Amersham Imager 600.

### qRT-PCR

Total RNA was isolated using the NucleoSpin RNA extraction kit (Macherey-Nagel), then reverse transcribed using the High-Capacity cDNA Reverse Transcription Kit (Applied Biosystems). qPCR of cDNAs was performed using Fast SYBR Green Master Mix (Applied Biosystems) on a QuantStudio 3 (ThermoFisher). RT-qPCR data were normalized to GAPDH or to G6PD as indicated in the figure legends. KicqStart primers to detect SIGIRR, IL1B, IL36RN, CXCL8, CXCL1, SDC4, IL18, IL18BP, IL36A, IL36B, IL36G, IL37, IL1F10, GAPDH, and G6PD were ordered from Sigma. Custom qRT-PCR primers for 16E6, 16E7, IL1A, IL1R1, IL1R2 were ordered from IDT with sequences as follows: 16E6 FWD: 5’ ATGGGAATCCATATGCTGTATGT, 16E6 REV: 5’ ACGGTTTGTTGTATTGCTGTTC, 16E7 FWD: 5’ AATGACAGCTCAGAGGAGGA, 16E7 REV: 5’ CGTAGAGTCACACTTGCAACA, IL1A FWD: 5’ CGCCAATGACTCAGAGGAAGA, IL1A REV: 5’ AGGGCGTCATTCAGGATGAA, IL1R1 FWD: 5’ AGAGGAAAACAAACCCACAAGG, IL1R1 REV: 5’ CTGGCCGGTGACATTACAGAT, IL1R2 FWD: 5’ TGGCACCTACGTCTGCACTACT, IL1R2 REV: 5’ TTGCGGGTATGAGATGAACG.

### RNAseq

Total RNA was isolated from three independent HFK cell populations +/- IL-1β treatment using the RNeasy mini kit (Qiagen). PolyA selection, reverse transcription, library construction, sequencing, and initial bioinformatic analysis were performed by Novogene. Differentially expressed genes were selected based on a 1.5- or 2.5-fold change and adjusted *p≤*0.05 cutoff or as noted. Gene ontology (GO) analysis to identify enriched biological processes was performed using DAVID analysis. RNA-seq data have been deposited in NCBI GEO with accession number GSE211730.

### Cancer genomic data analysis

Genomic mutation and tumor RNA-seq gene expression data from TCGA were analyzed using the cBioPortal.org graphical interface (21). RNA-seq V2 RSEM (RNA-Seq by Expectation Maximization) normalized expression values for individual genes were downloaded from cBioPortal.org. Statistical analysis of TCGA data and of PDX data was performed as in cBioPortal (21).

### Patient derived xenografts

The PDXs were previously established from surgical resections of treatment-naive HPV-positive OPSCC as described (71). Human tumors were engrafted subcutaneously in NSG mice and passaged at least twice before cryopreservation when they reached a volume of 0.5-1.0 cm^3^. Total tumor RNA was isolated using the QIAamp RNA Blood Mini Kit (Qiagen). All patient-derived materials and clinical data in this study were obtained from patients who underwent surgery to remove an oral cavity or oropharyngeal cancer. Patients were counseled preoperatively and provided informed consent under University of Pennsylvania IRB-approved protocol #417200 “Head and Neck Cancer Specimen Bank” (PI: D. Basu) by signing a combined informed consent and HIPAA form for use of tissue for research.

## ACKNOWLEDGEMENTS

We thank members of the White laboratory for helpful discussions related to these experiments. This research was supported by American Cancer Society grant 131661-RSG-18-048-01-MPC to EAW and by a pilot grant from the Penn Center for AIDS Research (CFAR), an NIH-funded program (P30 AI 045008). DB is supported by NIH/NIDCR R01 DE027185. SBDRC was funded by NIH grant P30 AR068589.

## AUTHOR CONTRIBUTIONS

Conception and design: PC, LKML, EAW. Acquisition of data: PC, HWK, LKML, EAW. Analysis and interpretation of data: PC, HWK, EAW. Drafting or revising the article: PC, DB, EAW. Contributing unpublished essential data or reagents: DB.

## FIGURE LEGENDS

**Supplemental Figure 1** | **High-risk oncoproteins dysregulate expression of genes involved in IL-1 signaling**. A) HFK were transduced with retroviruses encoding HPV16 E6 and HPV16 E7, HPV18 E6 and HPV18 E7, HPV6 E6 and HPV6 E7, or with matched vector control retroviruses and selected with appropriate antibiotics. RNA from the cell lines was analyzed by qRT-PCR using primers specific for IL1A, IL1B, and SIGIRR. Graphs display mean ± SD of the relative RNA level (vs G6PD), n=3 B) Protein lysates from the cell lines in (B) were separated by SDS-PAGE and analyzed in Western blots with antibodies to SIGIRR, pro-IL-1β, and Actin.

**Supplemental Figure 2** | **Somatic mutational burden in IL-1 pathway genes is comparable in HPV-negative and HPV-positive OPSCC**. Genomic mutation and copy number variation data from 28 HPV-negative and 53 HPV-positive OPSCC in The Cancer Genome Atlas (TCGA) was accessed using cBioPortal. Oncoprint displays specific genomic alterations in individual tumor samples.

**Supplemental Figure 3** | **Expression of IL-1 family cytokines and regulatory proteins in HPV-negative and HPV-positive patient-derived xenografts**. Total RNA was purified from 11 HPV-negative and 8 HPV-positive patient-derived xenografts (PDX) and qRT-PCR was used to assess gene expression of A) selected IL-1 family cytokines, B) IL1R1 (receptor), or C) IL1RN, IL18BP, and IL36RN (negative regulators). Data is expressed relative to G6PD expression (for IL18, IL1R1, and IL18BP) or to GAPDH expression (for IL36A, IL36B, IL36G, IL37, IL1F10, and IL36RN). Statistical significance was determined using unpaired t test with Welch’s correction. p-values below 0.1 are indicated. ns, not significant.

**Supplemental Figure 4** | **HPV16 oncoproteins suppress the transcriptional response to IL-1β**. Primary HFK stably expressing HPV16 E6 and HPV16 E7 or matched GFP control cells were treated with 0.5 ng/ml IL-1β for 3h or left untreated. RNA-seq and bioinformatic analysis was performed on polyA-selected RNA from triplicate cell populations. Heat map displays expression data for genes associated with GO term 0070555: Response to interleukin-1.

**Supplemental Table 1** | **Normalized qRT-PCR values for selected IL-1 pathway genes in HPV-positive and HPV-negative PDX**.

**Supplemental Table 2** | **List of differentially expressed genes (>1.5x up or down) in GFP/GFP cells +/- IL-1β**

**Supplemental Table 3** | **List of differentially expressed genes (>1.5x up or down) in 16E6/E7 cells +/- IL-1β**

**Supplemental Table 4** | **Pathway analysis of genes in four clusters shown in Figure 5A. DAVID analysis, Gene Ontology Biological process terms only**.

**Supplemental Table 5** | **Pathway analysis of genes in three subsets shown in Figure 5B. DAVID analysis, Gene Ontology Biological process terms only**.

**Supplemental Table 6** | **Plasmids used in the study**

## REFERENCES

1. Dyson N, Guida P, Munger K, Harlow E. 1992. Homologous sequences in adenovirus E1A and human papillomavirus E7 proteins mediate interaction with the same set of cellular proteins. J Virol 66:6893–902.

2. Dyson N, Howley PM, Munger K, Harlow E. 1989. The human papilloma virus-16 E7 oncoprotein is able to bind to the retinoblastoma gene product. Science 243:934–7.

3. Munger K, Werness BA, Dyson N, Phelps WC, Harlow E, Howley PM. 1989. Complex formation of human papillomavirus E7 proteins with the retinoblastoma tumor suppressor gene product. EMBO J 8:4099–105.

4. Scheffner M, Werness BA, Huibregtse JM, Levine AJ, Howley PM. 1990. The E6 oncoprotein encoded by human papillomavirus types 16 and 18 promotes the degradation of p53. Cell 63:1129–36.

5. Werness BA, Levine AJ, Howley PM. 1990. Association of human papillomavirus types 16 and 18 E6 proteins with p53. Science 248:76–9.

6. Hatterschide J, Bohidar AE, Grace M, Nulton TJ, Kim HW, Windle B, Morgan IM, Munger K, White EA. 2019. PTPN14 degradation by high-risk human papillomavirus E7 limits keratinocyte differentiation and contributes to HPV-mediated oncogenesis. Proc Natl Acad Sci U S A 116:7033–7042.

7. Hatterschide J, Brantly AC, Grace M, Munger K, White EA. 2020. A Conserved Amino Acid in the C Terminus of Human Papillomavirus E7 Mediates Binding to PTPN14 and Repression of Epithelial Differentiation. J Virol 94.

8. Hatterschide J, Castagnino P, Kim HW, Sperry SM, Montone KT, Basu D, White EA. 2022. YAP1 activation by human papillomavirus E7 promotes basal cell identity in squamous epithelia. Elife 11.

9. Johnson DE, Burtness B, Leemans CR, Lui VWY, Bauman JE, Grandis JR. 2020. Head and neck squamous cell carcinoma. Nat Rev Dis Primers 6:92.

10. Coussens LM, Werb Z. 2002. Inflammation and cancer. Nature 420:860–7.

11. Grivennikov SI, Greten FR, Karin M. 2010. Immunity, inflammation, and cancer. Cell 140:883–99.

12. Baker KJ, Houston A, Brint E. 2019. IL-1 Family Members in Cancer; Two Sides to Every Story. Front Immunol 10:1197.

13. Swindell WR, Beamer MA, Sarkar MK, Loftus S, Fullmer J, Xing X, Ward NL, Tsoi LC, Kahlenberg MJ, Liang Y, Gudjonsson JE. 2018. RNA-Seq Analysis of IL-1B and IL-36 Responses in Epidermal Keratinocytes Identifies a Shared MyD88-Dependent Gene Signature. Front Immunol 9:80.

14. Woodworth CD, Simpson S. 1993. Comparative lymphokine secretion by cultured normal human cervical keratinocytes, papillomavirus-immortalized, and carcinoma cell lines. Am J Pathol 142:1544–55.

15. Al-Sahaf S, Hunter KD, Bolt R, Ottewell PD, Murdoch C. 2019. The IL-1/IL-1R axis induces greater fibroblast-derived chemokine release in human papillomavirus-negative compared to positive oropharyngeal cancer. Int J Cancer 144:334–344.

16. Niebler M, Qian X, Hofler D, Kogosov V, Kaewprag J, Kaufmann AM, Ly R, Bohmer G, Zawatzky R, Rosl F, Rincon-Orozco B. 2013. Post-translational control of IL-1beta via the human papillomavirus type 16 E6 oncoprotein: a novel mechanism of innate immune escape mediated by the E3-ubiquitin ligase E6-AP and p53. PLoS Pathog 9:e1003536.

17. Ainouze M, Rochefort P, Parroche P, Roblot G, Tout I, Briat F, Zannetti C, Marotel M, Goutagny N, Auron P, Traverse-Glehen A, Lunel-Potencier A, Golfier F, Masson M, Robitaille A, Tommasino M, Carreira C, Walzer T, Henry T, Zanier K, Trave G, Hasan UA. 2018. Human papillomavirus type 16 antagonizes IRF6 regulation of IL-1beta. PLoS Pathog 14:e1007158.

18. Riva F, Bonavita E, Barbati E, Muzio M, Mantovani A, Garlanda C. 2012. TIR8/SIGIRR is an Interleukin-1 Receptor/Toll Like Receptor Family Member with Regulatory Functions in Inflammation and Immunity. Front Immunol 3:322.

19. Ndiaye C, Mena M, Alemany L, Arbyn M, Castellsagué X, Laporte L, Bosch FX, de Sanjosé S, Trottier H. 2014. HPV DNA, E6/E7 mRNA, and p16INK4a detection in head and neck cancers: a systematic review and meta-analysis. Lancet Oncol 15:1319–31.

20. Anonymous. 2015. Comprehensive genomic characterization of head and neck squamous cell carcinomas. Nature 517:576–82.

21. Cerami E, Gao J, Dogrusoz U, Gross BE, Sumer SO, Aksoy BA, Jacobsen A, Byrne CJ, Heuer ML, Larsson E, Antipin Y, Reva B, Goldberg AP, Sander C, Schultz N. 2012. The cBio cancer genomics portal: an open platform for exploring multidimensional cancer genomics data. Cancer Discov 2:401–4.

22. Gao J, Aksoy BA, Dogrusoz U, Dresdner G, Gross B, Sumer SO, Sun Y, Jacobsen A, Sinha R, Larsson E, Cerami E, Sander C, Schultz N. 2013. Integrative analysis of complex cancer genomics and clinical profiles using the cBioPortal. Sci Signal 6:pl1.

23. Mangino G, Chiantore MV, Iuliano M, Fiorucci G, Romeo G. 2016. Inflammatory microenvironment and human papillomavirus-induced carcinogenesis. Cytokine Growth Factor Rev 30:103–11.

24. Best SR, Niparko KJ, Pai SI. 2012. Biology of human papillomavirus infection and immune therapy for HPV-related head and neck cancers. Otolaryngol Clin North Am 45:807–22.

25. Hibma MH. 2012. The immune response to papillomavirus during infection persistence and regression. Open Virol J 6:241–8.

26. Hemmat N, Bannazadeh Baghi H. 2019. Association of human papillomavirus infection and inflammation in cervical cancer. Pathog Dis 77.

27. Georgescu SR, Mitran CI, Mitran MI, Caruntu C, Sarbu MI, Matei C, Nicolae I, Tocut SM, Popa MI, Tampa M. 2018. New Insights in the Pathogenesis of HPV Infection and the Associated Carcinogenic Processes: The Role of Chronic Inflammation and Oxidative Stress. J Immunol Res 2018:5315816.

28. Gameiro SF, Ghasemi F, Barrett JW, Koropatnick J, Nichols AC, Mymryk JS, Maleki Vareki S. 2018. Treatment-naive HPV+ head and neck cancers display a T-cell-inflamed phenotype distinct from their HPV-counterparts that has implications for immunotherapy. Oncoimmunology 7:e1498439.

29. Al-Sahaf S, Hendawi NB, Ollington B, Bolt R, Ottewell PD, Hunter KD, Murdoch C. 2021. Increased Abundance of Tumour-Associated Neutrophils in HPV-Negative Compared to HPV-Positive Oropharyngeal Squamous Cell Carcinoma Is Mediated by IL-1R Signalling. Front Oral Health 2:604565.

30. Liu X, Ma X, Lei Z, Feng H, Wang S, Cen X, Gao S, Jiang Y, Jiang J, Chen Q, Tang Y, Tang Y, Liang X. 2015. Chronic Inflammation-Related HPV: A Driving Force Speeds Oropharyngeal Carcinogenesis. PLoS One 10:e0133681.

31. Boccardo E, Lepique AP, Villa LL. 2010. The role of inflammation in HPV carcinogenesis. Carcinogenesis 31:1905–12.

32. Richards KH, Wasson CW, Watherston O, Doble R, Eric Blair G, Wittmann M, Macdonald A. 2015. The human papillomavirus (HPV) E7 protein antagonises an Imiquimod-induced inflammatory pathway in primary human keratinocytes. Sci Rep 5:12922.

33. Rm GDC, Bastos MM, Medeiros R, Oliveira PA. 2016. The NFκB Signaling Pathway in Papillomavirus-induced Lesions: Friend or Foe? Anticancer Res 36:2073–83.

34. Richards KH, Doble R, Wasson CW, Haider M, Blair GE, Wittmann M, Macdonald A. 2014. Human papillomavirus E7 oncoprotein increases production of the anti-inflammatory interleukin-18 binding protein in keratinocytes. J Virol 88:4173–9.

35. Abulkhir A, Samarani S, Amre D, Duval M, Haddad E, Sinnett D, Leclerc JM, Diorio C, Ahmad A. 2017. A protective role of IL-37 in cancer: a new hope for cancer patients. J Leukoc Biol 101:395–406.

36. Anselmo A, Riva F, Gentile S, Soldani C, Barbagallo M, Mazzon C, Feruglio F, Polentarutti N, Somma P, Carullo P, Angelini C, Bacci M, Mendolicchio GL, Voza A, Nebuloni M, Mantovani A, Garlanda C. 2016. Expression and function of IL-1R8 (TIR8/SIGIRR): a regulatory member of the IL-1 receptor family in platelets. Cardiovasc Res 111:373–84.

37. Cavalli G, Dinarello CA. 2018. Suppression of inflammation and acquired immunity by IL-37. Immunol Rev 281:179–190.

38. Dinarello CA, Nold-Petry C, Nold M, Fujita M, Li S, Kim S, Bufler P. 2016. Suppression of innate inflammation and immunity by interleukin-37. Eur J Immunol 46:1067–81.

39. Molgora M, Barajon I, Mantovani A, Garlanda C. 2016. Regulatory Role of IL-1R8 in Immunity and Disease. Front Immunol 7:149.

40. Nold-Petry CA, Lo CY, Rudloff I, Elgass KD, Li S, Gantier MP, Lotz-Havla AS, Gersting SW, Cho SX, Lao JC, Ellisdon AM, Rotter B, Azam T, Mangan NE, Rossello FJ, Whisstock JC, Bufler P, Garlanda C, Mantovani A, Dinarello CA, Nold MF. 2015. IL-37 requires the receptors IL-18Rα and IL-1R8 (SIGIRR) to carry out its multifaceted anti-inflammatory program upon innate signal transduction. Nat Immunol 16:354–65.

41. Wald D, Qin J, Zhao Z, Qian Y, Naramura M, Tian L, Towne J, Sims JE, Stark GR, Li X. 2003. SIGIRR, a negative regulator of Toll-like receptor-interleukin 1 receptor signaling. Nat Immunol 4:920–7.

42. Pontillo A, Bricher P, Leal VN, Lima S, Souza PR, Crovella S. 2016. Role of inflammasome genetics in susceptibility to HPV infection and cervical cancer development. J Med Virol 88:1646–51.

43. Goswami A, Bhuniya U, Chatterjee S, Mandal P. 2022. The influence of IL1RN VNTR polymorphism on HPV infection among some tribal communities. J Med Virol 94:752–760.

44. Tamandani DM, Sobti RC, Shekari M, Kaur S, Huria A. 2008. Impact of polymorphism in IL-1RA gene on the risk of cervical cancer. Arch Gynecol Obstet 277:527–33.

45. Qian N, Chen X, Han S, Qiang F, Jin G, Zhou X, Dong J, Wang X, Shen H, Hu Z. 2010. Circulating IL-1beta levels, polymorphisms of IL-1B, and risk of cervical cancer in Chinese women. J Cancer Res Clin Oncol 136:709–16.

46. Grulich AE, van Leeuwen MT, Falster MO, Vajdic CM. 2007. Incidence of cancers in people with HIV/AIDS compared with immunosuppressed transplant recipients: a meta-analysis. Lancet 370:59–67.

47. Angeletti PC, Zhang L, Wood C. 2008. The viral etiology of AIDS-associated malignancies. Adv Pharmacol 56:509–57.

48. Shiels MS, Cole SR, Kirk GD, Poole C. 2009. A meta-analysis of the incidence of non-AIDS cancers in HIV-infected individuals. J Acquir Immune Defic Syndr 52:611–22.

49. Dubrow R, Silverberg MJ, Park LS, Crothers K, Justice AC. 2012. HIV infection, aging, and immune function: implications for cancer risk and prevention. Curr Opin Oncol 24:506–16.

50. Robbins HA, Shiels MS, Pfeiffer RM, Engels EA. 2014. Epidemiologic contributions to recent cancer trends among HIV-infected people in the United States. Aids 28:881–90.

51. Wang CC, Silverberg MJ, Abrams DI. 2014. Non-AIDS-Defining Malignancies in the HIV-Infected Population. Curr Infect Dis Rep 16:406.

52. Coghill AE, Shiels MS, Suneja G, Engels EA. 2015. Elevated Cancer-Specific Mortality Among HIV-Infected Patients in the United States. J Clin Oncol 33:2376–83.

53. Silverberg MJ, Lau B, Achenbach CJ, Jing Y, Althoff KN, D’Souza G, Engels EA, Hessol NA, Brooks JT, Burchell AN, Gill MJ, Goedert JJ, Hogg R, Horberg MA, Kirk GD, Kitahata MM, Korthuis PT, Mathews WC, Mayor A, Modur SP, Napravnik S, Novak RM, Patel P, Rachlis AR, Sterling TR, Willig JH, Justice AC, Moore RD, Dubrow R. 2015. Cumulative Incidence of Cancer Among Persons With HIV in North America: A Cohort Study. Ann Intern Med 163:507–18.

54. Nittayananta W, Tao R, Jiang L, Peng Y, Huang Y. 2016. Oral innate immunity in HIV infection in HAART era. J Oral Pathol Med 45:3–8.

55. Park LS, Hernandez-Ramirez RU, Silverberg MJ, Crothers K, Dubrow R. 2016. Prevalence of non-HIV cancer risk factors in persons living with HIV/AIDS: a meta-analysis. Aids 30:273–91.

56. Coghill AE, Pfeiffer RM, Shiels MS, Engels EA. 2017. Excess Mortality among HIV-Infected Individuals with Cancer in the United States. Cancer Epidemiol Biomarkers Prev 26:1027–1033.

57. Hernandez-Ramirez RU, Shiels MS, Dubrow R, Engels EA. 2017. Cancer risk in HIV-infected people in the USA from 1996 to 2012: a population-based, registry-linkage study. Lancet HIV 4:e495–e504.

58. Vangipuram R, Tyring SK. 2019. AIDS-Associated Malignancies. Cancer Treat Res 177:1–21.

59. Lam JO, Bream JH, Sugar EA, Coles CL, Weber KM, Burk RD, Wiley DJ, Cranston RD, Reddy S, Margolick JB, Strickler HD, Wentz A, Jacobson L, Guo Y, Xiao W, Gillison ML, D’Souza G. 2016. Association of serum cytokines with oral HPV clearance. Cytokine 83:85–91.

60. Nixon DE, Landay AL. 2010. Biomarkers of immune dysfunction in HIV. Curr Opin HIV AIDS 5:498–503.

61. Stacey AR, Norris PJ, Qin L, Haygreen EA, Taylor E, Heitman J, Lebedeva M, DeCamp A, Li D, Grove D, Self SG, Borrow P. 2009. Induction of a striking systemic cytokine cascade prior to peak viremia in acute human immunodeficiency virus type 1 infection, in contrast to more modest and delayed responses in acute hepatitis B and C virus infections. J Virol 83:3719–33.

62. Bebell LM, Passmore JA, Williamson C, Mlisana K, Iriogbe I, van Loggerenberg F, Karim QA, Karim SA. 2008. Relationship between levels of inflammatory cytokines in the genital tract and CD4+ cell counts in women with acute HIV-1 infection. J Infect Dis 198:710–4.

63. Keating SM, Golub ET, Nowicki M, Young M, Anastos K, Crystal H, Cohen MH, Zhang J, Greenblatt RM, Desai S, Wu S, Landay AL, Gange SJ, Norris PJ. 2011. The effect of HIV infection and HAART on inflammatory biomarkers in a population-based cohort of women. Aids 25:1823–32.

64. Norris PJ, Pappalardo BL, Custer B, Spotts G, Hecht FM, Busch MP. 2006. Elevations in IL-10, TNF-alpha, and IFN-gamma from the earliest point of HIV Type 1 infection. AIDS Res Hum Retroviruses 22:757–62.

65. Palefsky J. 2006. Biology of HPV in HIV infection. Adv Dent Res 19:99–105.

66. von Sydow M, Sonnerborg A, Gaines H, Strannegard O. 1991. Interferon-alpha and tumor necrosis factor-alpha in serum of patients in various stages of HIV-1 infection. AIDS Res Hum Retroviruses 7:375–80.

67. White EA, Kramer RE, Tan MJ, Hayes SD, Harper JW, Howley PM. 2012. Comprehensive analysis of host cellular interactions with human papillomavirus E6 proteins identifies new E6 binding partners and reflects viral diversity. J Virol 86:13174–86.

68. White EA, Walther J, Javanbakht H, Howley PM. 2014. Genus beta human papillomavirus E6 proteins vary in their effects on the transactivation of p53 target genes. J Virol 88:8201–12.

69. Doench JG, Fusi N, Sullender M, Hegde M, Vaimberg EW, Donovan KF, Smith I, Tothova Z, Wilen C, Orchard R, Virgin HW, Listgarten J, Root DE. 2016. Optimized sgRNA design to maximize activity and minimize off-target effects of CRISPR-Cas9. Nat Biotechnol 34:184–191.

70. White EA, Sowa ME, Tan MJ, Jeudy S, Hayes SD, Santha S, Munger K, Harper JW, Howley PM. 2012. Systematic identification of interactions between host cell proteins and E7 oncoproteins from diverse human papillomaviruses. Proc Natl Acad Sci U S A 109:E260–7.

71. Facompre ND, Rajagopalan P, Sahu V, Pearson AT, Montone KT, James CD, Gleber-Netto FO, Weinstein GS, Jalaly J, Lin A, Rustgi AK, Nakagawa H, Califano JA, Pickering CR, White EA, Windle BE, Morgan IM, Cohen RB, Gimotty PA, Basu D. 2020. Identifying predictors of HPV-related head and neck squamous cell carcinoma progression and survival through patient-derived models. Int J Cancer 147:3236–3249.

